# The spread of resistance to imidacloprid is restricted by thermotolerance in natural populations of *Drosophila melanogaster*

**DOI:** 10.1101/511519

**Authors:** Alexandre Fournier-Level, Robert T Good, Stephen Wilcox, Rahul V Rane, Michelle Schiffer, Wei Chen, Paul Battlay, Trent Perry, Philip Batterham, Ary A Hoffmann, Charles Robin

**Author notes:** Corresponding authors: Alexandre Fournier-Level and Charles Robin.

## Abstract

Imidacloprid, the world’s most utilised insecticide^1^, has raised considerable controversy due to its harmful effects on non-pest species^2–6^ and there is increasing evidence showing that insecticides have become the primary selective force in many insect species^7–14^. The genetic response to insecticides is heterogeneous across population and environment^15–17^, leading to more complex patterns of genetic variation than previously thought. This motivated the investigation of imidacloprid resistance at different temperatures in natural populations of *Drosophila melanogaster* originating from four climate extremes replicated across two continents. Population and quantitative genomic analysis, supported by functional tests, demonstrated a polygenic basis to resistance and a major trade-off with thermotolerance. Reduced genetic differentiation at resistance-associated loci indicate enhanced gene flow at these loci. Resistance alleles showed stronger evidence of positive selection in temperate populations compared to tropical populations. Polygenic architecture and ecological factors should be considered when developing sustainable management strategies for both pest and beneficial insects.

## Introduction

Many organisms recurrently exposed to pesticides develop resistance, causing major issues for the sustainable control of pests and pathogens. Resistance emerges through many, if not any, mechanism that stop or limit the toxic compound, or toxic derivatives, from reaching its molecular target. However, understanding the extent to which ecological and evolutionary context shapes the adaptive response to insecticides remains challenging. While resistance emerging through mutations of major effect has been extensively described^18–20^, resistance that has a multifactorial basis has often been overlooked. Recent large-scale genomic projects underscore findings that the amplification of gene families often contributes to insecticide resistance^21–23^ and that, alongside new mutations, standing polygenic variation and ecological factors play an important role globally^24,25^. A quantitative basis to resistance is a theoretical requirement to support the many cases of recessive resistance, and also helps explain why resistance to new pesticides is occurring increasingly faster since the introduction of modern chemical controls (penicillin, DDT and 2,4-D for microbes, insects and weeds, respectively)^26–29^.

Under steady selection, alleles conferring resistance are expected to go to fixation and selective sweeps have indeed been widely observed for loci underlying resistance, one of the best examples being the *Cyp6g1* locus conferring resistance to DDT and many other chemicals in *Drosophila melanogaster*^*8,18*^. This locus has experienced a sequence of selective sweeps which increased the frequency of resistant alleles^30,31^. However, the ancestral susceptible allele was still evident in recent samples of natural genetic diversity, showing that highly favoured resistance alleles might not always fixate^32^. The persistence of such polymorphism suggests evolutionary trade-offs between resistance and other ecological factors^16,33^ which are poorly understood particularly for multifactorial resistance.

Resistance mutations often occur in genes that exert a primary, ancestral function (e.g the nervous system for insect or the photosystems for plants) while resistance is a secondary, derived function. Selective forces may act differently on resistance genes recruited from different pathways and induce a fitness cost for bearing the resistance allele and trade-offs may vary among populations depending on local adaptive pressures^34–36^. However, evidence of fitness costs associated with resistance are not that commonly reported in natural pest populations^37–40^. This could be due at least in part to a complex eco-evolutionary context that is difficult to reproduce in the laboratory and experimental measurements of fitness may not reflect those that are relevant in field conditions^41^. A way forward is to combine experiments that apply specific types of environmental variation with the use of population backgrounds that are specifically adapted to particular environmental conditions. The detection of trade-offs can then be facilitated by harnessing whole-genome sequencing approaches and a well characterised set of responses to experimental perturbations.

At the species range scale, populations evolve across multiple environments under different selection pressures^16^. This is particularly the case for widespread species including pests and disease vectors^42,43^. *D. melanogaster* is a cosmopolitan and synanthropic species that has a broad range and bears abundant diversity within natural populations^44^, ideal for testing complex evolutionary questions. Wild populations vary in their responses to temperature^45^ and insecticide^30^ with potential parallels across continents^46^ so it is now timely to investigate interactions between multiple genetic and environmental factors with population genomic resolution.

Here we explore how multiple field populations of *D. melanogaster* which have undergone long-term selection in different climates have dealt with the novel and ubiquitous short-term selection pressures associated with pesticides. Using a parallel sampling across the climate extremes of two continents, we build on the unmatched resources existing in *D. melanogaster* to comprehensively analyse the effect of Single Nucleotide Polymorphisms (SNPs), Insertion and Deletions (InDels), Transposable Elements (TEs), Copy-Number Variants (CNVs) and symbiotic micro-organisms on the genetic architectures of temperature and insecticide resistance. This provides a test of whether genetic differentiation observed for resistance genes interacts with thermotolerance at the phenotype and molecular level.

## Results

### Longevity in response to combined insecticide and temperature stress

In regions of extreme temperature and precipitation (Fig. 1A-B), 16 natural populations of *Drosophila melanogaster* were sampled outdoors in autumn with two collections per climate zone and per continent (Table S1). Individual flies (n=12,770) from 400 isofemale lines (i.e. the progeny of a single, mated field-collected female fly) were tested for longevity within 25 generations from collection, and were either exposed or not exposed to 1000ppm of imidacloprid diluted in sucrose media, at either 20°C or 30°C. Imidacloprid exposure reduced longevity by 83.9 hours (SE=4.9) producing a non-acute response while increasing the temperature from 20°C to 30°C reduced longevity by 118.1 hours (SE=4.1). The flies originating from hot climates had longer lifespan in the heat (ANOVA, *n*=7801, *F*_*1,7799*_=10.17, P=0.00143, Fig1C) consistent with anticipated climate adaptation, but had shorter lifespan when exposed to imidacloprid (ANOVA, *n*=10612, *F*_*1,10610*_=11.13, P=0.000854, Fig1C). The climate zone of origin as well as the continent of origin had a significant effect on longevity, however the models including population and isofemale line effects explained a greater proportion of variance (Table S2). 58% of the variance in time-to-death was explained by isofemale genotype effect and 32% by population effect, suggesting that genetic variation for longevity segregates more within population than among them. The responses to elevated temperature and imidacloprid computed from the effects of linear model with the best fit to the data were strongly negatively correlated, both at the population (*ρ*=-0.91, Fig. 1D) and at the isofemale genotype level (*ρ*=-0.68, Fig. 1E), revealing a trade-off between insecticide and elevated temperature tolerance within and among populations. The Australian populations were more resistant to imidacloprid than the American (Fig. 1D) with the exception of the population from Victoria (Australia).

**Figure 1:**
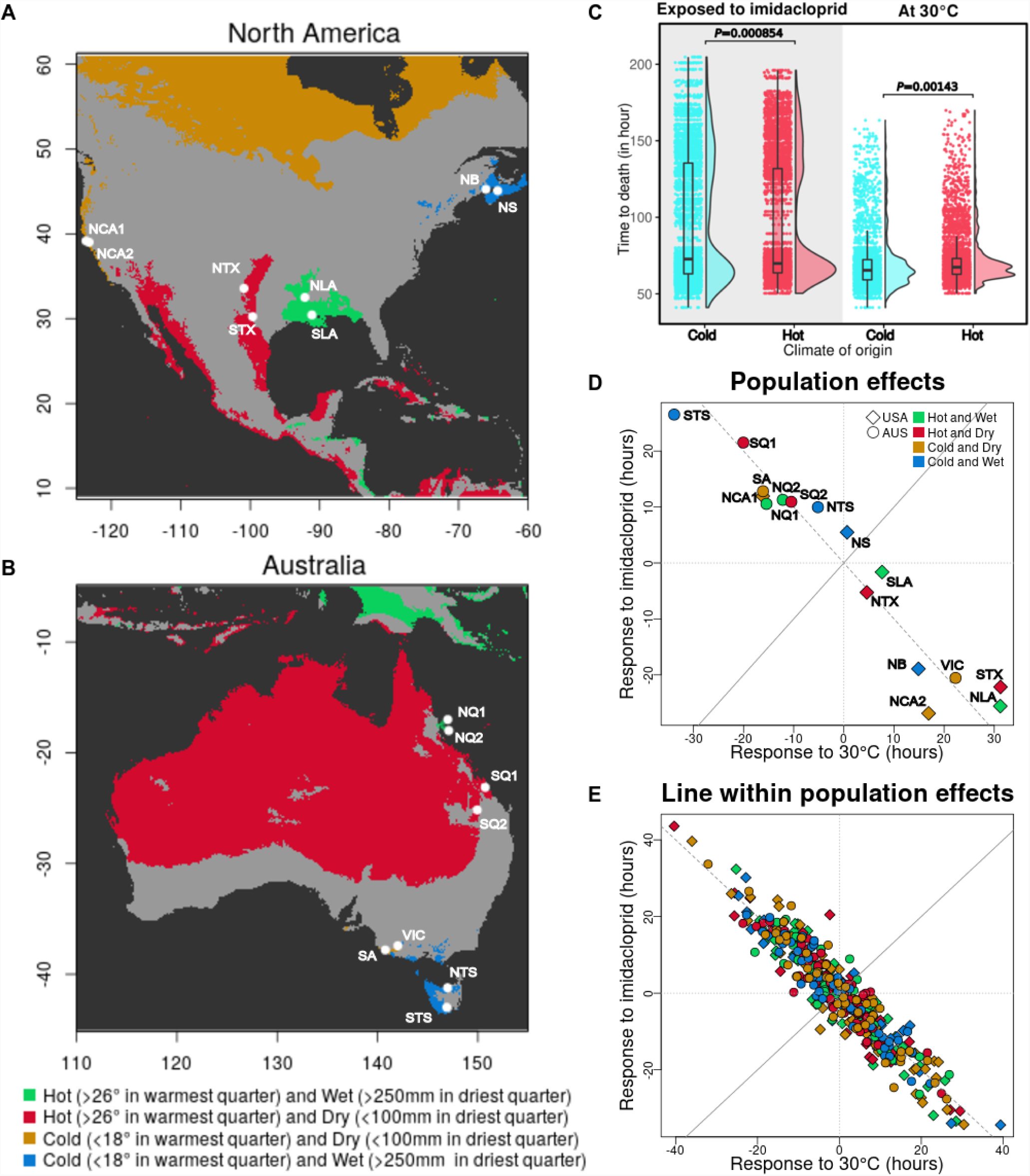
**A-B) Origin of the populations sampled and location of extreme climates in North America and Australia. C) Difference in the longevity of individual flies exposed to 1000ppm imidacloprid or to 30°C depending on their climate of origin. D-E) Mixed-linear model effects of population or isofemale line within population variation on longevity under elevated temperature** (30°C) **and insecticide stress** (1000ppm imidacloprid). Population labels are: NB: New Brunswick, NCA1: North California 1, NCA2: North California 2, NLA: North Louisiana, NQ1: North Queensland 1, NQ2: North Queensland 2, NS: Nova Scotia, NTS: North Tasmania, NTX: North Texas, SA: South Australia, SLA: South Louisiana, SQ1: South Queensland 1, SQ2: South Queensland 2, STS: South Tasmania, STX: South Texas, VIC: Victoria.

### Molecular variation associated with climate and resistance to imidacloprid

For each population at each temperature, every fly exposed to imidacloprid was assigned into one of five pools of resistance that covered the complete phenotypic distribution; each pool of genomic DNA was sequenced. The sequence reads were mapped to the *Drosophila melanogaster* “hologenome”, which includes the reference genome, and those of six of its most common microbes^47^ to call SNPs, TE insertions and CNV. SNP-based polymorphism and genetic diversity were consistent across populations (average *π*=0.0058 and *θ*_*Watterson*_=0.0059 for the autosomes, Table S2) with the exception of the North California 1 population that showed about two-fold more diversity (*π*=0.0116 and *θ*_*Watterson*_=0.0134). Genome-wide average estimates of *π* and *θ*_*Watterson*_ were not different (paired t-test, *n*=16, t=-1.746, P=0.101), suggesting populations were sampled at a demographic equilibrium without recent intense selective or founding events. Neither the temperature nor the precipitation at the location of origin was correlated with the level of polymorphism (GLM, *n*=16, P>0.15).

Using SNP alleles tagging cosmopolitan chromosomal inversions^48^, the *In(3R)Payne* inversion was found to be more prevalent in Australia and showed clinal variation from high frequency in North Queensland to low frequency in South Tasmania, consistent with previous findings^49^ (Table S3). The *In(2L)t* and *In(3R)Payne* inversions were more frequent in hot climates (three-way ANOVA testing continent, temperature and precipitation regime of origin, *n*=16, *In(2l)t: F*_*1,11*_=22.1, P=6.53×10^-4^ and *In(2l)Payne: F*_*1,11*_=22.0, P=6.63×10^-4^) and *In(3R)Mo* in wet climates (*F*_*1,11*_=4.973, P=0.0202). The only common microbe differentially present across populations was *Acetobacter pomorum* which was more frequent in North America (Table S4, *F*_*1,11*_=6.730, P=0.0249) and in cold climates in general (*F*_*1,11*_=5.643, P=0.0368). Interestingly, the presence of pathogenic bacteria *Pseudomonas entomophila* and *Gluconobacter morbifer* correlated with the increased lifespan of flies exposed to imidacloprid from North Tasmania and in North California 1 and South Australia (GLM, 5 pools, P=0.008, P=0.001 and P=0.006, respectively), always at 30°C. We also detected a positive effect of *Wolbachia pipientis* on longevity at 30°C in North Louisiana and North Texas (GLM, 5 pools, P=0.005, P=0.007).

Genome-Wide Association (GWA) between the SNPs and TEs and imidacloprid resistance revealed the influence of two major loci (Fig. 2A-B). These two loci were identified in 22 and 21 out of the 32 GWA tests (16 populations tested for two temperatures, Fig. S2), and were located within the coding sequence of the *Paramyosin* (*Prm*) gene and within the coding sequence plus 60kbp upstream of the *Nicotinic-Acetylcholine Receptor Alpha 3* (*nACHRα3*) gene, the latter encoding a subunit of the nicotinic-acetylcholine receptor, imidacloprid’s molecular target^50^. CRISPR deletion alleles of *nACHRα3* increased imidacloprid susceptibility while heterozygous MiMIC^51^ disruption for *Prm* increased susceptibility only at 20°C (Fig 3).

**Figure 2:**
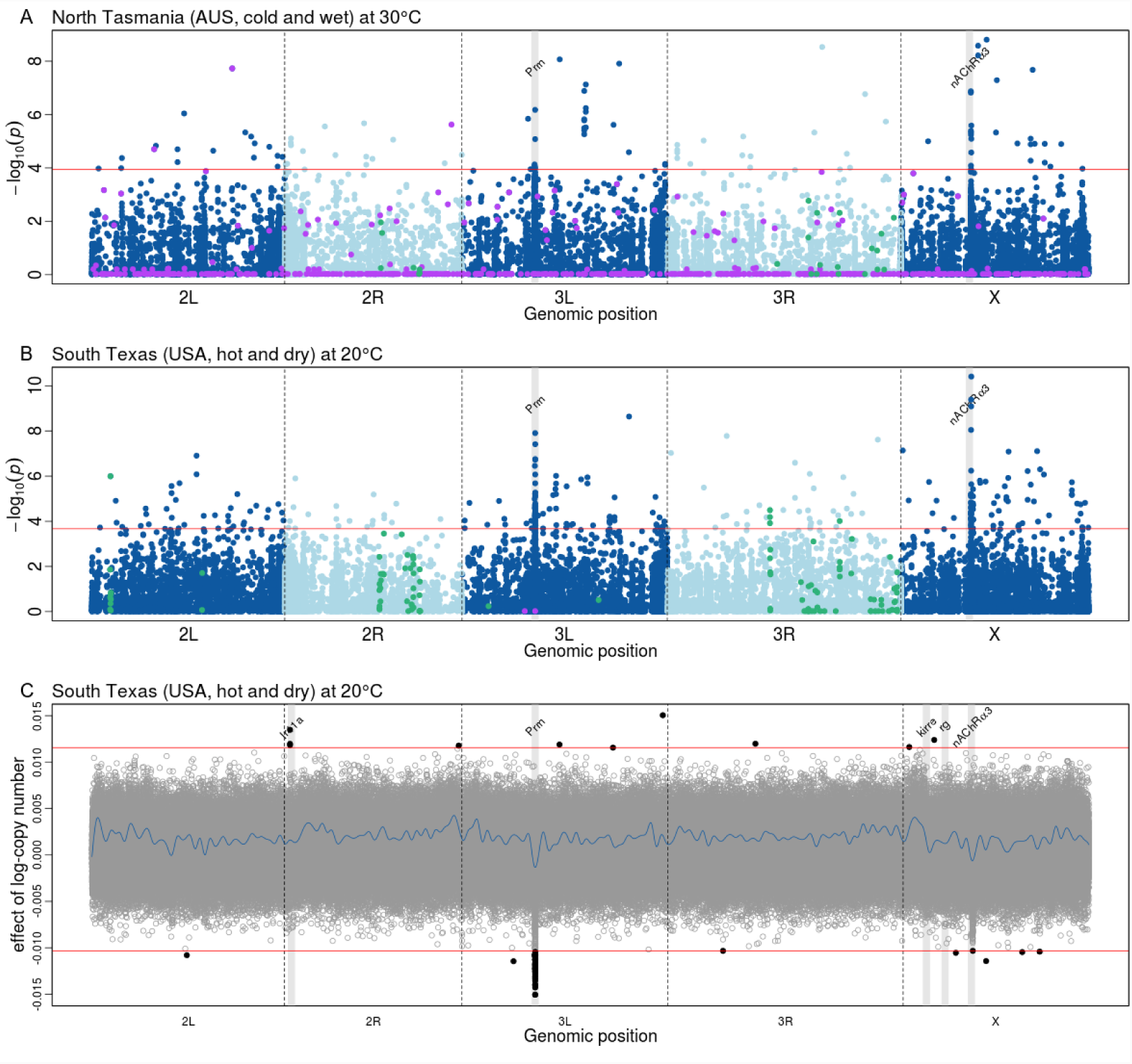
**A-B) Manhattan plots indicating the genomic position of polymorphisms associated with imidacloprid resistance.** Blue dots are SNP and small InDels, green dots are Inversion-tagging SNPs, purple dots are transposable elements and the red line indicates the FDR=5% threshold. Vertical shades of grey indicate the position of the two main candidate genes, *Paramyosin* and *N-Acetylcholine receptor α 3* located in position 3L:8,733,456-8,745,162 and X:8,277,227-8,438,674 of the *D. melanogaster* genome v6. **C) Effect of CNV on imidacloprid resistance by windows of 266bp.** Positive values indicate a greater number of copies increasing resistance. Grey and black dots are non-significant and significant CNV, respectively; the blue line represent a smoothed spline curve joining the individual dots (smoothing parameter λ=0.03); the red lines indicate the FDR=5% threshold. Vertical shades of grey indicate the position of the five main candidate genes: *Ionotropic receptor 41a, Paramyosin, Kin of Irre, Rugose* and *N-Acetylcholine receptor alpha 3*.

**Figure 3:**
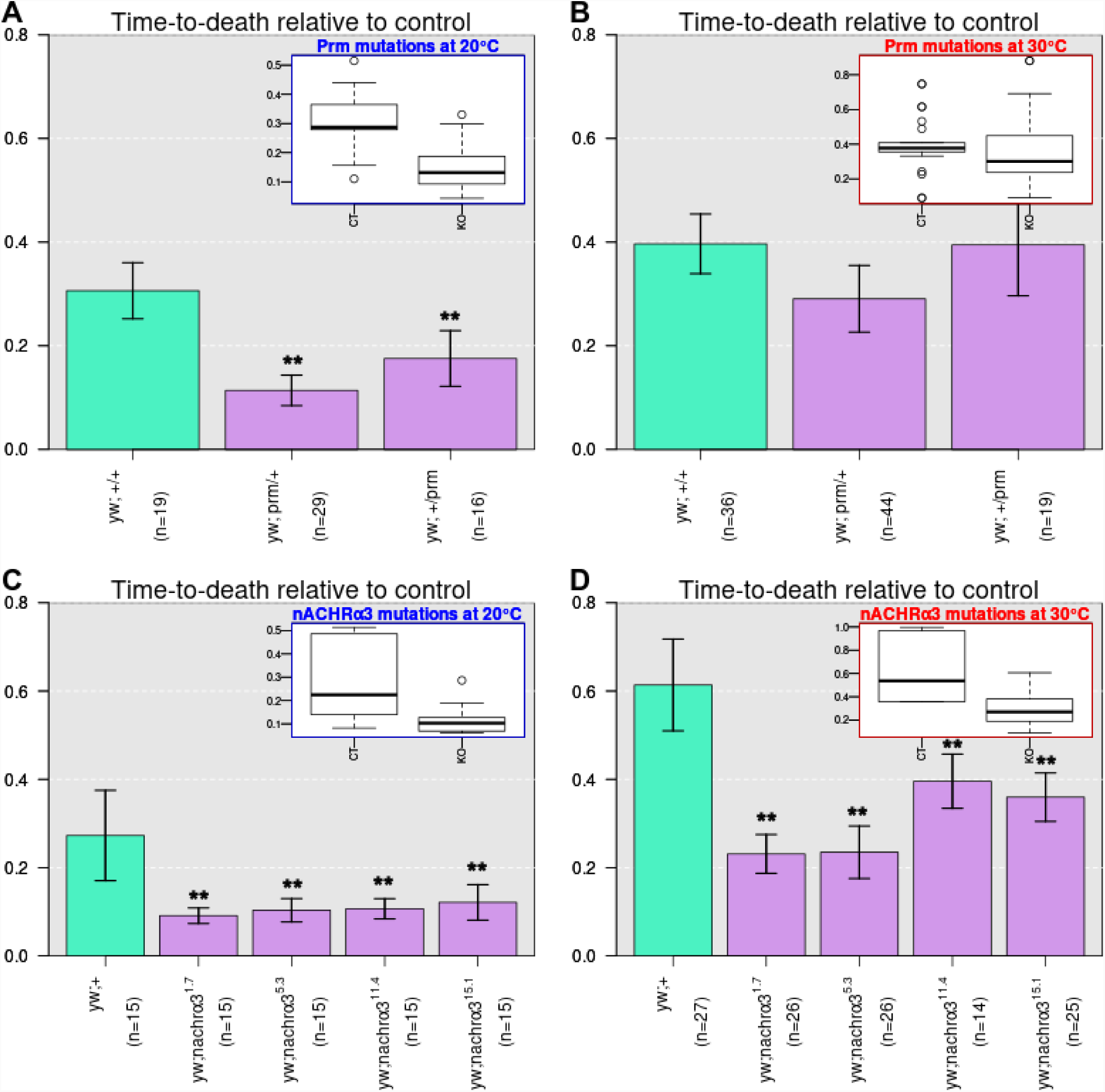
**A-D) Mean longevity +/- standard error of flies exposed to imidacloprid relative to control** (y-axis) **for wild type and mutants in candidate genes.** The top-right box in each panel is a scatter-boxplot recapitulating the difference in genetic effect between mutant and wildtype for each gene and temperature comparison in the same unit of relative longevity with respect to control; the thick shows the mean effect, the box shows the interquartile distribution and the whiskers, the 5%- and 95%-tile distribution. Test for *Prm* are shown in panel A and B, *nAChRα3* in panel C and D tested at 20**°**C (panel A and C) or 30**°**C (panel B and D). **: P-value<0.01 (t-test).

16 of the 26 TEs associated with imidacloprid resistance belonged to the R1A1-element family (analysis restricted to the Australian populations which have higher sequencing coverage overall). All but one R1A1-element located in position 2R:19,900,676-19,901,175 (*D. melanogaster* v6 assembly) in the North Tasmania population were rare with a frequency lower than 0.06, suggesting that in our data, TEs did not heavily contribute to the variation detected for insecticide resistance (Fig. S1). 62 inversion-tagging SNPs out of 473 were significantly associated with imidacloprid phenotypes in five American and seven Australian populations (Fig. S2) with 41 out of these 62 SNPs decreasing resistance to imidacloprid. 34% of all SNP tagging *In(2L)t* and 72% of all SNPs tagging *In(3R)Payne* showed a significant association in at least one GWA test which is higher than expected by chance. GWA tests between CNVs and imidacloprid resistance in the Texas populations further supported the presence of major effects at the two major loci detected through SNP-based GWA with fewer copies corresponding to increased imidacloprid resistance which is the opposite of knock-out mutation effects (Fig. 2C). However, the three CNVs most commonly influencing resistance across populations (in at least 10 out of 32 GWA tests, Fig. S3) all corresponded to higher copy-number increasing longevity. These CNVs located in position 2R:4,917,519-5,021,955, X:2,740,384-3,134,532 and X:5,085,759-5,254,864 colocalised with the *Ir41a*, the *kirre* and the *rg* genes, respectively.

### Patterns of population differentiation and selection among populations

The genetic differentiation (*F*_*st*_) between populations was correlated with geographic distance, consistent with isolation by distance (*r*_*Mantel*_=0.23, P<0.001) with no influence of latitude (distance from the equator, *r*_*Mantel*_=0.03, P=0.442). Overall less than half of the genetic differentiation was explained by climate distance (*r*_*partial Mantel*_=0.11, P=0.018). However, two regions at the top of chromosome 2L and bottom of chromosome 3R collocating with the *In(2L)t, In(3R)Mo* and *In(3R)Payne* inversions showed high genetic differentiation driven by climate consistent with previous findings^47,49,52^(Fig. 4A-B). Conversely, the genomic regions where candidate genes are located show little evidence of climate differentiation. We found that the average *F*_*st*_ value for genomic intervals containing resistance candidates identified across multiple GWA tests, was lower than expected (Monte Carlo simulation, P<0.006, Fig. 4D), highlighting minimal genetic differentiation for resistance candidate genes and supporting the hypothesis of an increased gene flow for these loci.

**Figure 4:**
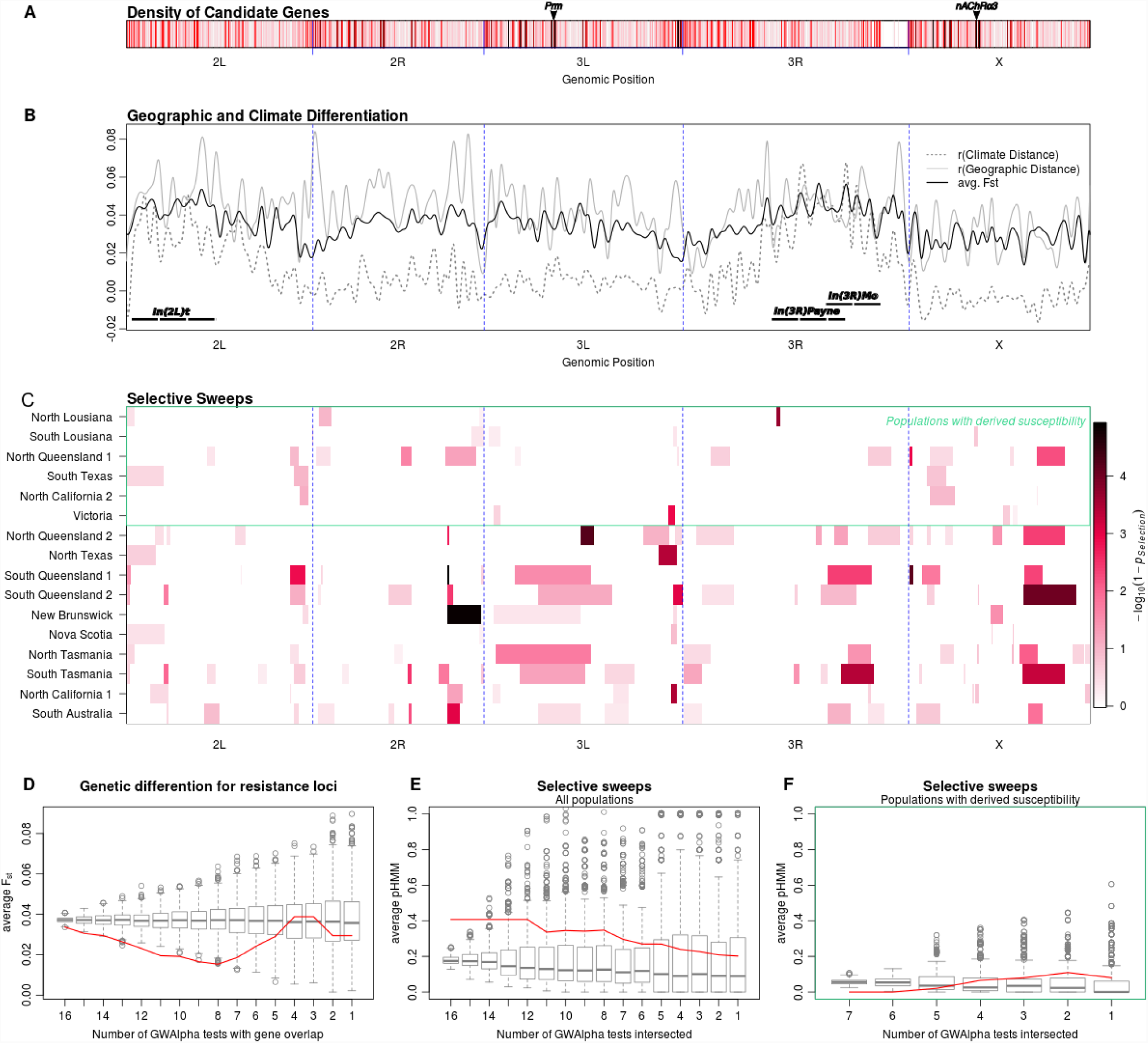
**A) Density of candidate genes along the genome**. Darker colours indicate higher density of candidate genes; The position of the two main canadidate gene *Prm* and *nAChRα3* is shown with an arrow **B) Average Fst and partial Mantel r statistic for correlations between Fst and geographic/climate distance over the genome** (sliding window of 5Kbp with smoothing parameter λ=0.05). The position of the three inversion associated with decayed resistance and increased climate adaptation *In(2L)t, In(3R)Mo* and *In(3R)Payne* is reported with long dashes. **C) Position of selective sweeps.** Selective sweeps were inferred based on the log-probability of transitioning from a neutral to a selected state (pHMM) inferred from a Hidden Markov Model based on site-frequency spectrum for each chromosome and each population. **D-F) Monte Carlo simulations testing the distribution of Fst among candidate genes** (solid red line) **compared to null genome-wide expectation** (box for interquartile and whiskers for inter-5%-ile distribution)**, the distribution of probabilities of selection in a Hidden Markov Model (pHMM) across all populations and the distribution of pHMM in populations with derived susceptibility**. The x-axis corresponds to the minimum number of GWA tests a given candidate gene was identified in (from the gene set found significantly associated in 16 or more GWA tests to the gene set found significantly associated in a one or more GWA test).

The evolutionary state of each allele, either ancestral or derived, was inferred by remapping the genome of the four most closely related species of *Drosophila* for which complete genomic sequences are available (*D. simulans, D. yakuba, D. erecta* and *D. sechellia*). For 23 out of 32 GWA tests, the resistance alleles predominantly corresponded to derived allele states while susceptible alleles were ancestral (χ^2^, maximum P<0.026 for these 23 GWAS). Three out of four tropical populations did not follow this pattern (both Louisiana and North Queensland 1 populations), with derived alleles conferring insecticide susceptibility at all temperatures (showed in the green box in Fig. 4C). This was also true for the South Texas, the North California 1 and the Victoria populations but only for the temperature furthest from their expected climatic preference (20°C, 30°C and 30°C, respectively). Using global freshwater pesticide data^53^, we did not find an effect of the level of pesticide pollution at the location of origin on insecticide resistance or the excess of derived allele conferring susceptibility. However, the excess of derived alleles conferring susceptibility positively affected the trade-off by increasing longevity at 30**°**C over longevity exposed to imidacloprid (GLM, P<0.0005).

Selective sweep analysis performed using Hidden Markov models of the allele frequency spectrum^54^ (Fig. 4C) strongly identified two regions in position 2R:20,779,995-20,989,995 and X:16,013,467-16,633,467 (top 1%-tile of selection probabilities averaged across populations) and a region in position 3L:23,339,400-23,849,400 (top 5%-tile) but did not include *Prm* or *nAChRα3*. *Para*, encoding the molecular target of organochloride and pyrethroid insecticides^19^, but never associated with resistance in natural populations of *D. melanogaster*, was located in the regions of chromosome X showing evidence of positive selection (top 1%-tile of selection probabilities). Selection probabilities among candidate genes identified in more than 13 out of 32 GWA tests were higher than expected (Monte Carlo simulation, P<0.005, Fig. 4E). The populations in which resistance alleles corresponded to ancestral states showed overall less evidence of positive selection compared to other populations (t-test, P=1.7×10^-9^). This pattern was accentuated for insecticide resistance loci: in the populations where the derived alleles increased susceptibility, among which three are tropical (green box in Fig4C), there was a lower than expected probability of selection (Monte Carlo simulation, P<0.001, Fig. 4F).

## Discussion

### Multifactorial bases to resistance

Resistance to imidacloprid in natural populations of *D. melanogaster* was shown to be controlled by a mixed genetic architecture involving major genes segregating across multiple populations and a polygenic background that exhibits a trade-off with thermotolerance. Of the two major genes we consistently identified, *nACHRα3* is functionally well characterised. It encodes one of the seven *α* subunit of the N-acetylcholine receptor which is the known target of imidacloprid^55^. *Prm* has been less extensively studied. In insects, it is known to encode important structural constituents of striated, obliquely striated and smooth muscles core thick filaments^56^. In *D. melanogaster, Prm* disruption was shown to affect indirect flight muscle stiffness and power generation^57^. Two studies have shown up-regulation of *Prm* in the context of insecticide resistance; in *Tribolium castaneum* exposed to insecticidal growth regulator diflubenzuron^58^ and in permethrin-resistant *Aedes aegyptii* after Zika virus infection^59^, respectively. The involvement of *Prm* could be a neuro-muscular compensation of a post-synaptic dysfunction as observed in human^60,61^, and in this case, it could act after the binding of imidacloprid or perhaps ameliorate the *nACHRα3 mutation.* The two major candidate genes from our study primarily point to target-site modes of resistance and this mechanism is underscored by significant enrichment of nervous system genes in the GWA analysis (Fig. S4, coupled with functional evidence for *ank* and *Pde9*, Fig. S5). Conversely, we only identified two genes belonging to the canonical detoxification genes family, and neither of these (*Cyp311a1* and *Cyp4s3*) are strong candidates for having roles in insecticide metabolism. The P450 *Cyp6g1*, that has previously been associated with imidacloprid metabolism^62–64^, was not detected in our analysis and inspection of a SNP tagging the CNV believed to be causal for the resistance (2R:12185332), is fixed in almost all Australian populations (the exception being a South Australian population) and was at high frequency in the north American populations we sampled (0.71 to 0.97). The analysis of microbial “hologenomes” adds another likely contributor to the multifactorial basis to variation in insecticide mortality. The microbes associated with resistance were all pathogenic, contrary to the climate- or continent-associated ones. This line of evidence leans toward a priming effect of immune defence mechanisms^65^ as opposed to the more common hypothesis of enhanced holometabolisation of toxic compounds being mediated through microorganisms^63,66–68^.

### Intra-genomic conflict could slow the evolution of resistance

We have probed multiple sources of molecular variation and showed that different classes of genomic change contribute to survivorship on insecticides and that they differ between temperatures. In particular, copy-number variation can be a major mechanism for resistance evolution^23,30^ and in this study, CNVs collocated with the two major resistance loci. While no significant pattern emerged through the analysis of transposable elements, cosmopolitan genomic inversions showed more consistent effects. *Payne* and *Mo*, the inversions associated with increasing susceptibility in our study, are likely to preserve gene complexes involved in climate adaptation^47,49^. These recombination-limiting genomic structures may also slow the pace of evolution, particularly when a novel selection pressure such as an insecticide interacts with a long-term one such as environmental stress. The fixation of a resistance allele can thus be considerably slowed down by the genomic context and result in the persistence of polymorphism irrespective of directional selection on the resistance allele. Nonetheless, this study also suggests that the genetic response to imidacloprid is conditional to the environment to which insects are pre-adapted. Insecticide resistance is thus fundamentally a complex trait likely to be both affected by genotype-by-environment interactions and by selection acting on a specific locus with different evolutionary potential. Due to a focus on variation segregating within populations, resistance mutations that have already reached fixation or that are near fixation (such as *Cyp6g1* alleles in Australian populations) do not arise in our genome wide association studies. However, a high fraction of resistance loci showed intermediate frequency with low levels of population differentiation relative to other variants at similar frequency across the genome. This suggests that gene flow is higher in these regions and that could be explained by selection favouring the spread of such variants. This is supported by the finding that imidacloprid resistance loci, notably those in temperate populations show signals of selection in their allele frequency spectrum. Given recessivity is typical, the increase in frequency of such polymorphisms foreshadow resistance outbreaks and motivates the identification of similar polymorphisms in pest species.

### The use of the eco-evolutionary context for managing resistance

Species that develop resistance to pesticides are typically widespread and they often are found to exhibit local adaptations that initially appear distinct from adaptations to control chemicals^69^. However here we have shown that the ecological genomic approach we have taken can identify important trade-offs between contrasting selective pressures. The peri-domestic ecology of the model insect *Drosophila melanogaster* was used as an environmental sentinel and the wealth of molecular resources available to understand its biology provides a powerful platform for testing multiple hypotheses. While it is well established that the same major insecticide resistance variants may arise independently in distantly related species (e.g. the same knockdown resistant amino acid mutations have occurred in many insect species^24^), it is less likely that genes contributing to the polygenic component will be common between species. However, the polygenes of different species are more likely to share molecular functions or act in common molecular pathways. Furthermore, the finding that the genomic architecture of insecticide resistance can point to evolutionary limits should encourage a systematic targeting of genomic regions where differentiation is high and/or where recombination is limited in pest and vector insects (many pest insects do indeed harbour chromosomal inversions). Here we identified such genomic regions by combining three experimental factors in the design of the study: rational sampling across climate extremes, traits being tested under multiple conditions and whole-genome sequencing. The population sampling ensured the ecological context of pre-adaptation to distinct temperature regimes was retained to its best extent and the tests at different temperatures led to physiological responses that increased the chance of identifying trade-offs. The pool-sequencing approach enabled efficient analysis of whole-genome data. The array of methods used in this study are readily transferable to most species inhabiting diverse environments, regardless of whether they are pests, disease vectors or beneficial species affected by pesticides.

### Conclusion

Knowledge of an insecticide-to-insect molecular interface alone, in the absence of a relevant environmental and genomic context, will likely not be enough to understand and predict resistance outbreaks, and will not provide an appreciation of the interplay of environment variation, genetic constraints and evolutionary outcomes. Ecological barriers seem capable of limiting the spread of functional genetic diversity, and such barriers need to be more systematically investigated across multiple populations and environments. Imidacloprid has been widely used in the last decades and has received considerable attention for its effect on beneficial insects in recent years. The response to imidacloprid in *D. melanogaster* is thus generally relevant even though the reductionist nature of the experiments would have overlooked other factors such as resistance to multiple other insecticides and climate variables which could add to the already complex situation described in our study. However, the evolutionary response it promotes is likely to be very similar to that of other classes of pesticide and the genomic mechanism of the response are also likely to be similar in other insects (inversion or CNV). Understanding resistance evolution across tiers of biological organisation including populations across the entire species range, genotypes within populations and explicit genomic context is thus becoming the new challenge. With such information, there is an opportunity to develop an improved understanding of the evolutionary trajectory of insecticide resistance in target pests and non-target species.

## Methods

### Fly collections, maintenance and bioassay

Natural fly populations were collected outdoors in October 2013 in North America and in March/April 2014 in Australia, corresponding to early Autumn for each continent (Fig. 1A-b and Table S1). 25 isofemale lines were established from each population and reared in the lab under constant light and temperature (25°C) conditions on Bloomington cornmeal media. Flies were kept for less than 25 generations in the lab before performing the experiments.

For each population, five 4 to 7 day old individual female flies from each isofemale line were individually exposed to an 1000ppm imidacloprid (Confidor®; Bayer Crop Science) diluted in 500mL of a 2% agarose/5% sucrose + 50mg/L tegosept media. Controls were exposed to the same mixture lacking imidacloprid. This design was tested at two different temperatures (20°C and 30°C) and replicated three times with flies from different generations to avoid batch effects. For each replicate, the time-to-death of 125 individual flies (5 flies per isofemale line) was recorded every 12 to 18 hours. For each replicate of each population the dead flies were immediately frozen and distributed into five pools (6.25%-, 24.5%-, 74.5%- and 94.75%-tile) based on their level of resistance as measured by time of death. This equates to 375 flies being distributed across 5 pools for each library synthesis (24, 72, 123, 72 and 24 flies respectively).

### Sample preparation and sequencing

Total genomic DNA was extracted using phenol/chloroform/isoamyl alcohol. Total DNA was quantified and 10ng was used to synthesise sequencing libraries using Illumina TrueSeq nano (Illumina, San Diego, USA) and barcode adapters kits (Agilent, Santa Clara, USA). Library quality and DNA concentration was measured on an Agilent 2200 TapeStation using D1000 DNA tapes (Agilent, Santa Clara, USA). The 160 libraries (16 populations x 2 temperatures x 5 times-to-death) corresponding to either the Australian or American samples were pooled and run on 7 lanes of the Illumina NextSeq 500 sequencer (Illumina, San Diego, USA) in 75bp paired-end reads mode.

### Genomic analysis

Paired-end sequences in FASTQ format were trimmed to remove adapter and barcodes and aligned to the *Drosophila melanogaster* “hologenome” consisting of the reference sequence v5.40 plus common microbes^47^ using the Burrows-Wheeler Aligner (BWA v0.7.12-r1039) with the *mem* option and default parameters^70^. PCR and optical duplicates were removed using Picard v1.112 and reads locally realigned around insertions/deletions using GATK v8.25. Alignment files in BAM format were merged for the different replicates using SAMtools and the resulting BAMs for the different pools of a given populations and temperature test were synchronised into MPILEUP using SAMtools and SYNC files using Popoolation2 v1.201. SNP and InDels were called using a custom script ensuring a Phred base quality score greater than q30, a minor allele frequency greater than 0.03, a base coverage greater than the 25%-tile of the corresponding pool. Finally, only SNPs present in both temperature tests (treated as biological replicates) were called. The criteria of presence in both temperature tests ensured a robust identification of polymorphisms.

TE were called using PopoolationTE2^71^ with the *D. melanogaster* v5.40 genome masked for the TEs present in the reference sequence and using a publicly available TE hierarchy for *D. melanogaster* (https://sourceforge.net/projects/popoolation-te2/files/walkthrough-refgenome.zip/download). As for the SNPs and InDels, the presence of a TE in a population was validated if detected for the two temperature tests (biological replicates) within a window of 5kbp. Chromosomal inversion diagnostic SNPs were retrieved from the literature^48^, the corresponding positions were retrieved directly from the MPILEUP file without further cutoffs and the relevant alleles were tested for association, but excluded from the other polymorphism and population analysis. CNV were called based on read-depth using the CNVkit software v0.9.3^72^. Frequency of symbionts in each pool was detected from the merged BAM alignment file by comparing the average read depth over a given symbiont genome divided the average read depth of the *D. melanogaster* genome using SAMtools and custom scripts.

### Statistical analysis

Mixed-linear models were computed using the lme4 package of the R software. Nested models were compared using an ANOVA procedure and compared using deviance and Akaike information criteria (Table S2). The best model was:

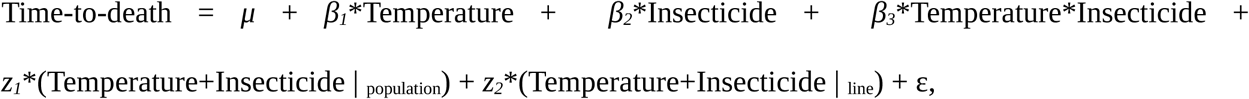

where μ is the intercept, *β*’s are the fixed effects of increasing the temperature from 20 to 30°C, the fixed effects of increasing the imidacloprid exposure from 0 to 1000ppm in the and the effect of their interaction, *z*’s are the random effects of increasing the temperature or the insecticide exposure among populations and among lines within population and ε the residual term.

Pool-based genome-wide association tests were performed with the GWAlpha package^73^ using maximum likelihood estimation and significance was calculated using empirical p-values with a Benjamini and Hochenberg correction for multiple testing for a false discovery rate of FDR=5% on the loci whose coverage was within the interquartile range for a given pool. Candidate genes were identified based on the excessive 95% proportion of associated SNPs or InDels in the neighbouring 5kbp window using the D. melanogaster genome annotation v5.57. Network and enrichment analysis were performed using the R spider pipeline^74^ under the model D2 (one missing connection allowed) using 2047 gene name entries for D. melanogaster from the KEGG and Reactome databases (1^st^ of December 2017). 225 gene identifiers out of 745 were recognised and 178 had a pathway information entry in the databases.

The Monte Carlo simulations used to test population genetic hypothesis were conducted by randomly shifting the circularised genomic position a 1000 times. The targeted summary statistic was then computed for the 1000 permutations to draw the expected null distribution and compare it to the observed statistic to derive the reported p-value.

### Population genetics analysis

For all population genetics analysis, each pool alignment file was down-sampled to even coverage for the 10 pool BAM alignment files for a given population (5 resistance pools x 2 temperatures) and merged into a single BAM file. Estimates for *π* and *θ* where obtained from the down-sampled merged BAM files using the Npstats package^75^ for windows of 5kbp only retaining polymorphism located in positions with quality phred > q30. Pairwise *F*_*st*_ between each population were estimated using the same setup as for *π* and *θ* using the Popoolation2 package^76^. Mantel and Partial Mantel tests were performed using the R\vegan package using 1000 permutations for p-value calculation. Climate distance of 0 corresponded to populations sharing both similar temperature and precipitation at their location of origin, 0.5 when sharing only temperature or precipitation or 1 otherwise. Ancestral and derived states for each SNP and InDel polymorphism were determined by “shredding” the reference genomes for four closely related *Drosophila* species (*D. simulans, D. yakuba, D. erecta* and *D. sechellia*) using the Wgsim script (https://github.com/lh3/wgsim) generating 2.5M 500bp-long fragments for each species that were re-mapped onto the *D. melanogaster* v5.40 genome using BWA. Polymorphic sites were then called using the *call* function from the BCFtools package v1.3.1 (https://samtools.github.io/bcftools). For each SNP/InDel position, the ancestral state was assigned if three out of five (including *D. melanogaster* reference genome) states were identical or left undetermined otherwise. Selective sweeps were detected based on probability of selection in a Hidden Markov Model^54^ implemented in the Pool-hmm package v1.4.3^77^. Allele frequency spectrum was computed for each chromosome using a random 10% of the sites that had a coverage greater than 10X from the down-sampled merged BAM file for each population. We performed a sensitivity analysis to determine the transition probability *k* from neutral to selected state and determined that for k>10e^-4^ the set selective sweeps detected remained consistent, so k was set to 10e^-4^.

All custom codes are deposited on GitHub (https://github.com/aflevel/IMIresist_vs_TEMPtol) and the derived data are deposited on figshare (DOI:<u>10.26188/5b592f305226b</u>). Raw sequencing data generated in this study are available upon request.

### Functional tests

Fly stocks were ordered at the Bloomington Drosophila Stock Center and the crossing design used to obtain the tested lines are reported in Table S7. Deletions of *nAChRα3* were generated using two small guide (sg)RNAs flanking the targeted region designed using CRISPR Optimal Target Finder (http://tools.flycrispr.molbio.wisc.edu/targetFinder/), ActinCas9 gDNA was checked to ensure no SNPs existed and then the nAChRα3 Del sgRNA 1 (GTCGCCGCAAACGCACACGA) and nAChRα3 Del sgRNA2 (GCCCATCGATATAAAGCTAT) were ordered as oligonucleotides and each ligated separately into pU6-BbsI-gRNA plasmids as per the published protocol (http://flycrispr.molbio.wisc.edu/protocols/gRNA). These were co-injected (each plasmid at 200ng/μl in water) into ActinCas9 flies following a standard microinjection protocol. Given the *nAChRα3* gene is located on chromosome X, injected adult male survivors were initially crossed back to AC9 and F1 males collected. A wing was clipped from males, DNA extracted and diagnostic PCR (Primers F 5’GTGTGACTGTATTGGTGCTG3’ and R 5’CACACACAGTCTGATGGAGC3’) for deletion events performed. Flies with products smaller than the expected size (3590kb) were individually crossed to virgin female FM7 balancer flies and these were backcrossed to remove the FM7 balancer, producing the four independent homozygous deletion strains of *nAChRα3* used in this study (1.7, 5.3, 11.4 and 15.1). Off-target analysis is reported in section 4.1 of the Supplementary Data.

## Acknowledgements

Thanks to David Begun, Sean Myles and Craig Hart for sharing their laboratory facilities; Philippa Griffin for fly collection, Kristy Charles at Foursights Wines, David Herbert at Herbert Vineyard, Peter Dixon at Henty Estate, Jonathan Leahy at Becker’s Vineyard, John Seago at Ponchartrain Vineyard and Landry Vineyards for providing access to their estate. This work was supported by the Human Frontier in Sciences Long Term fellowship #LT000907/2012-L awarded to AFL.

## Author contributions

AFL and CR designed the study and wrote the manuscript with contribution from TP and AAH; AFL, RTG and SW performed the experiments; AFL analysed the data with contribution from AAH, RVR and Paul Battlay; AAH, Paul Battlay and TP revised the manuscript; AAH, MS, Phil Batterham, TP and WC provided new genetic material.

## References

1. Cressey, D. The bitter battle over the world’s most popular insecticides. Nature 551, 156–158 (2017).

2. Cernansky, R. Controversial pesticides found in honey samples from six continents. Nature (2017). doi:10.1038/nature.2017.2276.

3. Sureda Anfres, M. Controversial insecticides linked to wild bee declines. Nature (2016). doi:10.1038/nature.2016.2044.

4. Rundlöf, M. et al. Seed coating with a neonicotinoid insecticide negatively affects wild bees. Nature 521, 77–80 (2015).

5. Hallmann, C. A., Foppen, R. P. B., van Turnhout, C. A. M., de Kroon, H. & Jongejans, E. Declines in insectivorous birds are associated with high neonicotinoid concentrations. Nature 511, 341–343 (2014).

6. Henry, M. et al. A Common Pesticide Decreases Foraging Success and Survival in Honey Bees. Science (80-.). 336, 348–350 (2012).

7. Schmidt, J. M. et al. Insights into DDT resistance from the drosophila melanogaster genetic reference panel. Genetics 207, (2017).

8. Battlay, P., Schmidt, J. M., Fournier-Level, A. & Robin, C. Genomic and Transcriptomic Associations Identify a New Insecticide Resistance Phenotype for the Selective Sweep at the Cyp6g1 Locus of Drosophila melanogaster. G3 Genes|Genomes|Genetics g3.116.031054 (2016). doi:10.1534/G3.116.03105.

9. Miles, A. et al. Genetic diversity of the African malaria vector Anopheles gambiae. Nature 552, 96 (2017).

10. Lynd, A. et al. Field, Genetic, and Modeling Approaches Show Strong Positive Selection Acting upon an Insecticide Resistance Mutation in Anopheles gambiae s.s. Mol. Biol. Evol. 27, 1117–1125 (2010).

11. Barnes, K. G. et al. Genomic Footprints of Selective Sweeps from Metabolic Resistance to Pyrethroids in African Malaria Vectors Are Driven by Scale up of Insecticide-Based Vector Control. PLoS Genet. 13, e1006539 (2017).

12. Jones, C. M. et al. Footprints of positive selection associated with a mutation (N1575Y) in the voltage-gated sodium channel of Anopheles gambiae. doi:10.1073/pnas.120147510.

13. Zamoum, T. et al. Does insecticide resistance alone account for the low genetic variability of asexually reproducing populations of the peach-potato aphid Myzus persicae? Heredity (Edinb). 94, 630–639 (2005).

14. Anderson, C. J. et al. Hybridization and gene flow in the mega-pest lineage of moth, Helicoverpa. Proc. Natl. Acad. Sci. U. S. A. 115, 5034–5039 (2018).

15. Woodcock, B. A. et al. Country-specific effects of neonicotinoid pesticides on honey bees and wild bees. Science (80-.). 356, 1393–1395 (2017).

16. Fournier-Level, A. et al. Behavioural response to combined insecticide and temperature stress in natural populations of Drosophila melanogaster. J. Evol. Biol. 29, (2016).

17. Glunt, K. D., Oliver, S. V., Hunt, R. H. & Paaijmans, K. P. The impact of temperature on insecticide toxicity against the malaria vectors Anopheles arabiensis and Anopheles funestus. Malar. J. 17, 131 (2018).

18. Daborn, P. J. et al. A single p450 allele associated with insecticide resistance in Drosophila. Science 297, 2253–6 (2002).

19. Pittendrigh, B., Reenan, R., ffrench-Constant, R. H. & Ganetzky, B. Point mutations in the Drosophila sodium channel gene para associated with resistance to DDT and pyrethroid insecticides. Mol. Gen. Genet. 256, 602–10 (1997).

20. Fournier, D., Bride, J. M., Hoffmann, F. & Karch, F. Acetylcholinesterase. Two types of modifications confer resistance to insecticide. J. Biol. Chem. 267, 14270–4 (1992).

21. Traverso, L. et al. Comparative and functional triatomine genomics reveals reductions and expansions in insecticide resistance-related gene families. PLoS Negl. Trop. Dis. 11, e0005313 (2017).

22. Faucon, F. et al. Identifying genomic changes associated with insecticide resistance in the dengue mosquito Aedes aegypti by deep targeted sequencing. Genome Res. 25, 1347–1359 (2015).

23. Faucon, F. et al. In the hunt for genomic markers of metabolic resistance to pyrethroids in the mosquito Aedes aegypti: An integrated next-generation sequencing approach. PLoS Negl. Trop. Dis. 11, e0005526 (2017).

24. Ffrench-Constant, R. H. The molecular genetics of insecticide resistance. Genetics 194, 807–15 (2013).

25. Messer, P. W. & Petrov, D. A. Population genomics of rapid adaptation by soft selective sweeps. Trends Ecol. Evol. 28, 659–69 (2013).

26. Ventola, C. L. The Antibiotic Resistance Crisis: Part 1: Causes and Threats. Pharm. Ther. 40, 277–283 (2015).

27. Tabashnik, B. E. & Carrière, Y. Surge in insect resistance to transgenic crops and prospects for sustainability. Nature Biotechnology 35, 926–935 (2017).

28. Mallet, J. The evolution of insecticide resistance: Have the insects won? Trends in Ecology and Evolution 4, 336–340 (1989).

29. Munro, A. Economics and Biological Evolution 1. Environ. Resour. Econ. 9, 429–449 (1997).

30. Schmidt, J. M. et al. Copy Number Variation and Transposable Elements Feature in Recent, Ongoing Adaptation at the <italic>Cyp6g1</italic> Locus. PLoS Genet 6, e1000998 (2010).

31. Garud, N. R. et al. Recent Selective Sweeps in North American Drosophila melanogaster Show Signatures of Soft Sweeps. PLOS Genet. 11, e1005004 (2015).

32. Schmidt, J. M. et al. Insights into DDT Resistance from the Drosophila melanogaster Genetic Reference Panel. Genetics 207, 1181–1193 (2017).

33. Labbé, P. et al. Forty years of erratic insecticide resistance evolution in the mosquito Culex pipiens. PLoS Genet. 3, e205 (2007).

34. Andersson, D. I. & Hughes, D. Antibiotic resistance and its cost: is it possible to reverse resistance? Nat. Rev. Microbiol. 8, 260 (2010).

35. Kliot, A. & Ghanim, M. Fitness costs associated with insecticide resistance. Pest Manag. Sci. 68, 1431–7 (2012).

36. Vila-Aiub, M. M., Neve, P. & Powles, S. B. Fitness costs associated with evolved herbicide resistance alleles in plants. New Phytol. 184, 751–767 (2009).

37. ffrench-Constant, R. H. & Bass, C. Does resistance really carry a fitness cost? Current Opinion in Insect Science 21, 39–46 (2017).

38. Wu, C., Davis, A. S. & Tranel, P. J. Limited fitness costs of herbicide-resistance traits in Amaranthus tuberculatus facilitate resistance evolution. Pest Manag. Sci. in press, (2017).

39. Melnyk, A. H., Wong, A. & Kassen, R. The fitness costs of antibiotic resistance mutations. Evol. Appl. 8, 273–283 (2015).

40. Bourguet, D., Guillemaud, T., Chevillon, C. & Raymond, M. Fitness costs of insecticide resistance in natural breeding sites of the mosquito Culex pipiens. Evolution 58, 128–35 (2004).

41. Cousens, R. D. & Fournier-Level, A. Herbicide resistance costs: what are we actually measuring and why? Pest Manag. Sci. (2017). doi:10.1002/ps.481.

42. Nkya, T. E. et al. Insecticide resistance mechanisms associated with different environments in the malaria vector Anopheles gambiae: a case study in Tanzania. Malar. J. 13, 28 (2014).

43. Sternberg, E. D. & Thomas, M. B. Local adaptation to temperature and the implications for vector-borne diseases. Trends Parasitol. 30, 115–122 (2014).

44. Lack, J. B., Lange, J. D., Tang, A. D., Corbett-Detig, R. B. & Pool, J. E. A Thousand Fly Genomes: An Expanded Drosophila Genome Nexus. Mol. Biol. Evol. 33, 3308–3313 (2016).

45. Hoffmann, A. A., Anderson, A. & Hallas, R. Opposing clines for high and low temperature resistance in Drosophila melanogaster. Ecol. Lett. 5, 614–618 (2002).

46. Turner, T. L., Levine, M. T., Eckert, M. L. & Begun, D. J. Genomic analysis of adaptive differentiation in Drosophila melanogaster. Genetics 179, 455–73 (2008).

47. Kapun, M., Fabian, D. K., Goudet, J. & Flatt, T. Genomic Evidence for Adaptive Inversion Clines in Drosophila melanogaster. Mol. Biol. Evol. 33, 1317–1336 (2016).

48. Kapun, M., Van Schalkwyk, H., McAllister, B., Flatt, T. & Schlötterer, C. Inference of chromosomal inversion dynamics from Pool-Seq data in natural and laboratory populations of Drosophila melanogaster. Mol. Ecol. 23, 1813–1827 (2014).

49. Rane, R. V., Rako, L., Kapun, M., Lee, S. F. & Hoffmann, A. A. Genomic evidence for role of inversion 3RP of Drosophila melanogaster in facilitating climate change adaptation. Mol. Ecol. 24, 2423–2432 (2015).

50. Matsuda, K. et al. Neonicotinoids: insecticides acting on insect nicotinic acetylcholine receptors. Trends Pharmacol. Sci. 22, 573–580 (2001).

51. Venken, K. J. T. et al. MiMIC: a highly versatile transposon insertion resource for engineering Drosophila melanogaster genes. Nat. Methods 8, 737–43 (2011).

52. Lavington, E. & Kern, A. D. The Effect of Common Inversion Polymorphisms In(2L)t and In(3R)Mo on Patterns of Transcriptional Variation in Drosophila melanogaster. G3 (Bethesda). 7, 3659–3668 (2017).

53. Vörösmarty, C. J. et al. Global threats to human water security and river biodiversity. Nature 467, 555–561 (2010).

54. Boitard, S., Schlotterer, C., Nolte, V., Pandey, R. V. & Futschik, A. Detecting Selective Sweeps from Pooled Next-Generation Sequencing Samples. Mol. Biol. Evol. 29, 2177–2186 (2012).

55. Buckingham, S. D., Lapied, B., Le Corronc, H., Grolleau, F. & Sattelle, D. B. Imidacloprid actions on insect neuronal acetylcholine receptors. J. Exp. Biol. 200, 2685–2692 (1997).

56. Cervera, M., Arredondo, J. J. & Ferreres, R. M. in Nature’s Versatile Engine: Insect Flight Muscle Inside and Out 76–85 (Springer US, 2006). doi:10.1007/0-387-31213-7_.

57. Liu, H. et al. Paramyosin phosphorylation site disruption affects indirect flight muscle stiffness and power generation in Drosophila melanogaster. Proc. Natl. Acad. Sci. U. S. A. 102, 10522–7 (2005).

58. Merzendorfer, H. et al. Genomic and proteomic studies on the effects of the insect growth regulator diflubenzuron in the model beetle species Tribolium castaneum. Insect Biochem. Mol. Biol. 42, 264–276 (2012).

59. Zhao, L., Alto, B., Shin, D. & Yu, F. The Effect of Permethrin Resistance on Aedes aegypti Transcriptome Following Ingestion of Zika Virus Infected Blood. Viruses 10, 470 (2018).

60. Gonzalez-Freire, M., de Cabo, R., Studenski, S. A. & Ferrucci, L. The Neuromuscular Junction: Aging at the Crossroad between Nerves and Muscle. Front. Aging Neurosci. 6, 208 (2s014).

61. Takamori, M. Synaptic Homeostasis and Its Immunological Disturbance in Neuromuscular Junction Disorders. Int. J. Mol. Sci. 18, (2017).

62. Joussen, N., Heckel, D. G., Haas, M., Schuphan, I. & Schmidt, B. Metabolism of imidacloprid and DDT by P450 CYP6G1 expressed in cell cultures of Nicotiana tabacum suggests detoxification of these insecticides in Cyp6g1-overexpressing strains of Drosophila melanogaster, leading to resistance. Pest Manag. Sci. 64, 65–73 (2008).

63. Fusetto, R., Denecke, S., Perry, T., O’Hair, R. A. J. & Batterham, P. Partitioning the roles of CYP6G1 and gut microbes in the metabolism of the insecticide imidacloprid in Drosophila melanogaster. Sci. Rep. 7, 11339 (2017).

64. Denecke, S. et al. Multiple P450s and Variation in Neuronal Genes Underpins the Response to the Insecticide Imidacloprid in a Population of Drosophila melanogaster. Sci. Rep. 7, (2017).

65. Broderick, N. A., Raffa, K. F. & Handelsman, J. Midgut bacteria required for Bacillus thuringiensis insecticidal activity. Proc. Natl. Acad. Sci. U. S. A. 103, 15196–9 (2006).

66. Cheng, D. et al. Gut symbiont enhances insecticide resistance in a significant pest, the oriental fruit fly Bactrocera dorsalis (Hendel). Microbiome 5, 13 (2017).

67. Dada, N., Sheth, M., Liebman, K., Pinto, J. & Lenhart, A. Whole metagenome sequencing reveals links between mosquito microbiota and insecticide resistance in malaria vectors. Sci. Rep. 8, 2084 (2018).

68. Kikuchi, Y. et al. Symbiont-mediated insecticide resistance. Proc. Natl. Acad. Sci. U. S. A. 109, 8618–22 (2012).

69. Hoffmann, A. A. Rapid adaptation of invertebrate pests to climatic stress? Curr. Opin. Insect Sci. 21, 7–13 (2017).

70. Li, H. Aligning sequence reads, clone sequences and assembly contigs with BWA-MEM. (2013). at <http://arxiv.org/abs/1303.3997>

71. Kofler, R., Gómez-Sánchez, D. & Schlötterer, C. PoPoolationTE2: Comparative Population Genomics of Transposable Elements Using Pool-Seq. Mol. Biol. Evol. 33, 2759–2764 (2016).

72. Talevich, E., Shain, A. H., Botton, T. & Bastian, B. C. CNVkit: Genome-Wide Copy Number Detection and Visualization from Targeted DNA Sequencing. PLOS Comput. Biol. 12, e1004873 (2016).

73. Fournier-Level, A., Robin, C. & Balding, D. J. GWAlpha: genome-wide estimation of additive effects (alpha) based on trait quantile distribution from pool-sequencing experiments. Bioinformatics btw805, btw805 (2016).

74. Antonov, A. V., Schmidt, E. E., Dietmann, S., Krestyaninova, M. & Hermjakob, H. R spider: A network-based analysis of gene lists by combining signaling and metabolic pathways from Reactome and KEGG databases. Nucleic Acids Res. 38, W78–W83 (2010).

75. Ferretti, L., Ramos-Onsins, S. E. & Pérez-Enciso, M. Population genomics from pool sequencing. Mol. Ecol. 22, 5561–5576 (2013).

76. Kofler, R., Pandey, R. V. & Schlötterer, C. PoPoolation2: identifying differentiation between populations using sequencing of pooled DNA samples (Pool-Seq). Bioinformatics 27, 3435–6 (2011).

77. Boitard, S. et al. Pool-hmm: a Python program for estimating the allele frequency spectrum and detecting selective sweeps from next generation sequencing of pooled samples. Mol. Ecol. Resour. 13, 337–40 (2013).

