## Supplementary Information for "The spread of resistance to imidacloprid is restricted by thermotolerance in natural populations of *Drosophila melanogaster*"

### 1) Population sampling

#### *1.1 Identification of of sampling locations of extreme climates*

Sampling zone was restricted to North America and Australia and locations with an altitude lower than 700m. Hot regions were defined using the Worldclim data (www.worldclim.org) as locations where the mean temperature during the warmest quarter of the year (BIO10) was greater than 26°C (or lesser than 18°C for cold regions); Wet regions were defined as locations where the minimum precipitation during the driest quarter of the year (BIO17) was greater than 250mm (or lesser than 100mm for dry regions).

#### *1.2 Fly collection*

Fly populations were primarily collected in vineyards (North California 1 and 2, North Louisiana, South Texas, Nova Scotia, North Tasmania, South Australia and Victoria) or orchards (North Queensland 1 and 2, North Texas and South Queensland 1) with the exception of the South Louisiana, New Brunswick, South Queensland 1 and South Tasmania which were trapped using buckets with ripped bananas as baits in open environments.

| Climate | Population | Continent | Location | Coordinates in ° | | Temperature in °C during the warmest quarter | Precipitation in mm during the driest quarter | Collection date | Min. and max number of generation from Collection to assay |
|---|---|---|---|---|---|---|---|---|---|
| | | | | Latitude | Longitude | | | | |
| Hot and Wet | North Lousiana | N. America | West Monroe | 32.5 | -92.14 | 27.1 | 245 | 9/2013 | 18 - 24 |
| | South Lousiana | N. America | Baton Rouge | 30.43 | -91.12 | 26.9 | 315 | 9/2013 | 8 - 24 |
| | North Queensland 1 | Australia | Innisfail Nth | -17.48 | 146 | 26.6 | 263 | 2/2014 | 8 - 23 |
| | North Queensland 2 | Australia | Innisfail Sth | -17.51 | 145.98 | 26.6 | 258 | 2/2014 | 5 - 10 |
| Hot and Dry | North Texas | N. America | Lubbock | 33.57 | -100.9 | 26.5 | 57 | 9/2013 | 8 - 24 |
| | South Texas | N. America | Stonewall | 30.23 | -99.66 | 26.4 | 94 | 9/2013 | 3 - 16 |
| | South Queensland 1 | Australia | Isla Gorge | -25.17 | 149.93 | 25.7 | 97 | 3/2014 | 14 - 20 |
| | South Queensland 2 | Australia | Yeppoon | -23.13 | 150.73 | 26.2 | 97 | 5/2014 | 2 - 8 |
| Cold and Wet | New Brunswick | N. America | St John | 45.25 | -66.07 | 16 | 283 | 10/2013 | 15 - 21 |
| | Nova Scotia | N. America | Gaspereau | 45.08 | -64.37 | 17.1 | 267 | 10/2013 | 5 - 7 |
| | North Tasmania | Australia | Lucaston | -41.23 | 146.98 | 16.7 | 136 | 4/2014 | 4 - 6 |
| | South Tasmania | Australia | Hillwood | -43 | 146.9 | 14.4 | 168 | 4/2014 | 2 - 9 |
| Cold and Dry | North California 1 | N. America | Anderson Valley Nth | 39.15 | -123.65 | 15.4 | 19 | 10/2013 | 9 - 20 |
| | North California 2 | N. America | Anderson Valley Sth | 39 | -123.21 | 18.6 | 15 | 10/2013 | 18 - 24 |
| | South Australia | Australia | Mt Gambier | -37.82 | 140.77 | 17.7 | 80 | 4/2014 | 14 - 17 |
| | Victoria | Australia | Hamilton | -37.43 | 142.02 | 18 | 93 | 5/2014 | 2 – 4 |

**Table S1:** Sampling design of the *Drosophila melanogaster* natural populations from extreme climates

### 2) Linear modelling

The initial null model was set as the model including the effect of temperature (df=1) and imidacloprid exposure (df=1) and their interaction (df=3) as fixed effects. Nested models were tested by including the effect of temperature and imidacloprid exposure and their interaction nested in the random effect of the population or line within population as random effects:

0- Time-to-death = $\mu + \beta_1$*Temperature + $\beta_2$*Insecticide + ε

1- Time-to-death = $\mu + \beta_1$*Temperature + $\beta_2$*Insecticide + $z_1$*(Temperature+Insecticide | $_{\text{population}}$) + ε

2- Time-to-death = $\mu + \beta_1$*Temperature + $\beta_2$*Insecticide + $z_1$*(Temperature+Insecticide | $_{\text{population}}$) + $z_2$*(Temperature+Insecticide | $_{\text{line}}$) + ε

3- Time-to-death = $\mu + \beta_1$*Temperature + $\beta_2$*Insecticide + $\beta_3$*Temperature*Insecticide + $z_1$*(Temperature+Insecticide | $_{\text{population}}$) + $z_2$*(Temperature+Insecticide | $_{\text{line}}$) + ε,

where μ is the intercept, $\beta$'s are the fixed effects of increasing the temperature from 20 to 30°C, the fixed effects of increasing the imidacloprid exposure from 0 to 1000ppm in the and the effect of their interaction, $z$'s are the random effects of increasing the temperature or the insecticide exposure among populations and among lines within population and ε the residual term.

Optimal model was selected based on minimum deviance and AIC as presented in Table S2.

| | Model Selection | | | | Proportion of variance explained | | |
|---|---|---|---|---|---|---|---|
| | Resid. Df | Resid. Dev | dAIC | weight | Line | Population | Total |
| Null | 12766 | 23751928.6 | 2992.1 | 0 | NA | NA | NA |
| Climate zone | 12759 | 132216 | 2831.3 | | NA | NA | 0.406 |
| Continent | 12759 | 132013.1 | 2628.5 | | NA | NA | 0.439 |
| Continent + Climate zone | 12753 | 131863.9 | 2487.1 | | NA | NA | 0.522 |
| Population | 12759 | 131028.8 | 1641.6 | 0 | NA | 0.521 | 0.521 |
| Population + Line | 12753 | 130118.2 | 743.2 | 0 | 0.378 | 0.313 | 0.691 |
| Population + Line with temperature x insecticide interaction | 12745 | 129361.6 | 0 | 1 | 0.319 | 0.581 | 0.9 |

**Table S2:** Comparison of linear-mixed models used to explain time-to-death.

Goodness of fit of the model and distribution of the residuals were plotted to validate the assumption of homoscedasticity and normal distribution of the residuals (Figure S1).

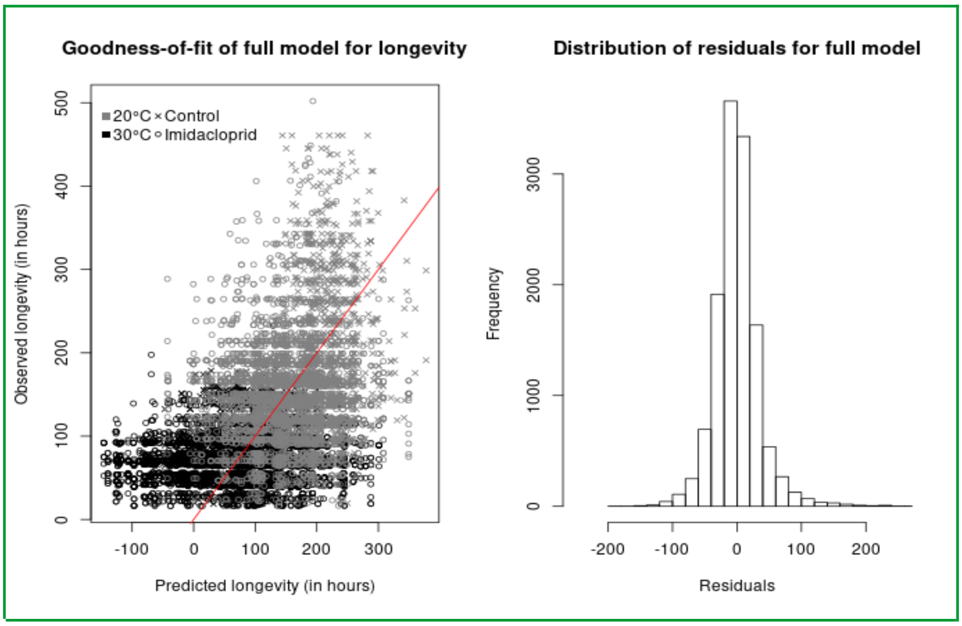


**Figure S1:** Predicted Time-to-death in model 3 compared to observed values (left), red line indicating perfect fit and distribution of residuals for model 3 (right)**.**

### 3) Sequencing output and polymorphism detection

#### *3.1 Sequencing strategy*

The targeted sequencing output was 1200Mb of sequence data for each of the 160 libraries (16 populations x 2 temperature x 5 levels of resistance). This was achieved using 3.5 Illumina NextSeq500/550 v2 kits and pooling together all 80 independently barcoded libraries (American and Australian populations kept apart due to limited number of barcodes) and adjusting the relative concentration of each library for each run until the target was reached for every library. 69.3 to 86.7% of the sequence reads mapped to the *D. melanogaster* reference genome v5.40 plus microbes. Based on the alignment file after removing PCR duplicates and performing local realignment around indels, the mean coverage for the euchromatic DNA regions (chromosome 2L, 2R, 3L, 3R, 4 and X) within each pool was 18.3X, ranging from a minimum of 7.6X to a maximum of 26.3X per library (Table S3).

#### *3.2 Calling Single Nucleotide Polymorphims, Transposable Elements and Copy Number Variants*

The higher sequencing coverage of the Australian populations resulted in greater power to detect TE insertions and higher specificity to detect CNV, therefore more TEs and less CNV were identified in these populations compared to the North American.

| Population | Sequencing output | | Molecular Diversity and Polymorphism | | | | |
|---|---|---|---|---|---|---|---|
| | Total mapped reads (75bp) count | Average Coverage per pool | $\pi$* | $\theta_{Watterson}$* | # SNP | # TE | # CNV |
| North Lousiana | 214670735 | 11.3X | 0.0063 (0.0042) | 0.0064 (0.0043) | 125499 | 40 | 58 |
| South Lousiana | 594528502 | 12.9X | 0.0044 (0.0019) | 0.0044 (0.002) | 131394 | 212 | 90 |
| North Queensland 1 | 297276933 | 24.2X | 0.0071 (0.0049) | 0.0076 (0.0054) | 244853 | 1187 | 11 |
| North Queensland 2 | 350716072 | 22.5X | 0.0066 (0.0044) | 0.0068 (0.0045) | 143376 | 646 | 14 |
| North Texas | 277390394 | 18.3X | 0.0059 (0.004) | 0.0059 (0.004) | 87560 | 229 | 33 |
| South Texas | 566375446 | 14.5X | 0.006 (0.0042) | 0.006 (0.0042) | 122969 | 64 | 37 |
| South Queensland 1 | 505985522 | 23.4X | 0.0065 (0.0043) | 0.0067 (0.0044) | 107536 | 2105 | 10 |
| South Queensland 2 | 527014680 | 25.7X | 0.0064 (0.0043) | 0.0066 (0.0044) | 124427 | 1325 | 27 |
| New Brunswick | 232205244 | 12.2X | 0.0055 (0.0037) | 0.0055 (0.0037) | 105086 | 36 | 68 |
| Nova Scotia | 144083076 | 15.7X | 0.0055 (0.0039) | 0.0055 (0.0039) | 122137 | 40 | 82 |
| North Tasmania | 244871869 | 26.3X | 0.0056 (0.0039) | 0.0057 (0.004) | 86828 | 1633 | 6 |
| South Tasmania | 583437144 | 24.9X | 0.0053 (0.0036) | 0.0054 (0.0036) | 93451 | 256 | 3 |
| North California 1 | 228422727 | 12.1X | 0.0116 (0.011) | 0.0134 (0.0129) | 94344 | 134 | 122 |
| North California 2 | 507927490 | 7.6X | 0.0048 (0.0036) | 0.0048 (0.0036) | 97102 | 33 | 34 |
| South Australia | 549281580 | 22.5X | 0.0058 (0.004) | 0.0059 (0.0041) | 129750 | 647 | 7 |
| Victoria | 404120591 | 18.1X | 0.0055 (0.0038) | 0.0055 (0.0038) | 103653 | 172 | 8 |

**Table S3:** Summary statistics for sequencing output and levels of polymorphism. *: average value calculated for euchromatic autosomes with the average value for the X chromosome reported between parenthesis.

*3.3 Calling chromosomal inversions*

| Population | Frequency of cosmopolitan inversions | | | | | | |
|---|---|---|---|---|---|---|---|
| | In(2L)t | In(2R)Ns | In(3L)P | In(3R)K | In(3R) Payne | In(3R)C | In(3R)Mo |
| North Lousiana | 0.25 | 0.08 | 0 | 0.02 | 0.15 | 0.01 | 0.06 |
| South Lousiana | 0.05 | 0.13 | 0.03 | 0.37 | 0.08 | 0.01 | 0 |
| North Queensland 1 | 0.3 | 0.07 | 0.15 | 0.01 | 0.7 | 0.01 | 0 |
| North Queensland 2 | 0.27 | 0.11 | 0.09 | 0 | 0.66 | 0.01 | 0 |
| North Texas | 0.41 | 0.08 | 0 | 0 | 0.1 | 0 | 0.03 |
| South Texas | 0.38 | 0.04 | 0.01 | 0 | 0.15 | 0.01 | 0.01 |
| South Queensland 1 | 0.21 | 0.09 | 0.06 | 0 | 0.26 | 0.03 | 0 |
| South Queensland 2 | 0.14 | 0.04 | 0 | 0 | 0.44 | 0 | 0 |
| New Brunswick | 0.06 | 0 | 0.01 | 0 | 0 | 0 | 0.15 |
| Nova Scotia | 0.07 | 0.03 | 0 | 0 | 0 | 0.02 | 0.11 |
| North Tasmania | 0.02 | 0.01 | 0 | 0 | 0.01 | 0 | 0 |
| South Tasmania | 0 | 0.02 | 0 | 0 | 0.06 | 0 | 0.01 |
| North California 1 | 0.01 | 0 | 0 | 0.04 | 0 | 0.02 | 0.03 |
| North California 2 | 0.02 | 0 | 0 | 0 | 0.02 | 0.01 | 0.03 |
| South Australia | 0.05 | 0.06 | 0 | 0.02 | 0 | 0.03 | 0 |
| Victoria | 0.06 | 0.08 | 0 | 0 | 0 | 0.01 | 0.01 |

**Table S4:** Frequency of chromosomal inversion across populations.

*3.3 Detection of microbes and endosymbionts*

| Population | Avg. Ratio of microbial to D. melanogaster DNA in samples | | | | | | | |
|---|---|---|---|---|---|---|---|---|
| | A. pomorum | G. morbifer | P. alcali-faciens | P. burhodo-granariea | P. entomo-phila | P. rettgeri | S. cerevisiae | W. pipientis |
| North Lousiana | 0.0017 | 0.0001 | 0 | 0.002 | 0 | 0.0001 | 0.0019 | 0.3763 |
| South Lousiana | 0.0086 | 0.0002 | 0 | 0.0015 | 0 | 0.0001 | 0.0015 | 0.6281 |
| North Queensland 1 | 0.0065 | 0.0001 | 0 | 0.0016 | 0 | 0.0001 | 0.0012 | 0.3363 |
| North Queensland 2 | 0.0069 | 0.0001 | 0 | 0.0017 | 0 | 0.0001 | 0.0014 | 0.4608 |
| North Texas | 0.0078 | 0.0001 | 0 | 0.0019 | 0 | 0.0002 | 0.0015 | 0.376 |
| South Texas | 0.0043 | 0.0003 | 0 | 0.0013 | 0 | 0.0002 | 0.0015 | 1.0475 |
| South Queensland 1 | 0.0045 | 0.0001 | 0 | 0.0017 | 0.0001 | 0.0001 | 0.0018 | 0.6604 |
| South Queensland 2 | 0.0026 | 0.0001 | 0 | 0.002 | 0 | 0.0001 | 0.002 | 0.6975 |
| New Brunswick | 0.0122 | 0 | 0 | 0.0016 | 0 | 0.0001 | 0.0013 | 0.3994 |
| Nova Scotia | 0.0132 | 0.0001 | 0.0001 | 0.0027 | 0 | 0.0001 | 0.0021 | 0.5172 |
| North Tasmania | 0.0089 | 0.0001 | 0 | 0.002 | 0 | 0.0001 | 0.0016 | 0.401 |
| South Tasmania | 0.0029 | 0.0001 | 0 | 0.0018 | 0 | 0.0001 | 0.002 | 0.6316 |
| North California 1 | 0.0088 | 0.0001 | 0.0002 | 0.0012 | 0 | 0.0001 | 0.0018 | 0.6348 |
| North California 2 | 0.0145 | 0.0001 | 0 | 0.002 | 0 | 0.0001 | 0.0019 | 0.7118 |
| South Australia | 0.0041 | 0.0001 | 0.0001 | 0.002 | 0 | 0.0001 | 0.0019 | 0.6634 |

**Table S5:** Frequency of microbes relative to *D. melanogaster* among populations.

### 4) Genome-wide Association tests

*4.1- Association between imidacloprid resistance and SNPs, TEs and chromosomal inversions*

**Figure S2: Manhattan plots indicating the genomic position of polymorphisms associated with imidacloprid resistance.** Blue dots are SNP and small InDels, green dots are Inversion-tagging SNPs, purple dots are transposable elements and the red line indicates the FDR=5% threshold. Vertical shades of grey indicate the position of the two main candidate genes, Paramyosin and N-Acetylcholine receptor *α* 3 located in position 3L:8,733,456-8,745,162 and X:8,277,227-8,438,674 of the *D. melanogaster* genome v6, respectively.

*(Figure S2 on next page)*

*4.2- Association between imidacloprid resistance and CNVs*

**Figure S3: Effect of CNV on imidacloprid resistance by windows of 266bp.** Positive values indicate a greater number of copies increasing resistance. Grey and black dots are non-significant and significant CNV, respectively; the blue line represent a smoothed spline curve joining the individual dots (smoothing parameter $\lambda$=0.03)**;** the red line indicates the FDR=5% threshold. Vertical shades of grey indicate the position of the five main candidate genes: *Ionotropic receptor 41a*, *Paramyosin*, *Kin of Irre*, *Rugose* and *N-Acetylcholine receptor α 3*.

*(Figure S3 on next page)*

### 4) Functional tests

#### *4.1 Functional validation for major effect genes*

*Paramyosyn* and *N-Acetylcholine receptor α 3* were the two major loci identified in the most consistently across GWAS (Fig. 2a-b).
For the *nAChRα3* validation involving the use of novel CRISPR deletion strains, off-target regions for the sgRNA used to generate the four homozygous delection strains of *N-Acetylcholine receptor α 3* were identified using Fly CRISPR Optimal Target Finder (http://tools.flycrispr.molbio.wisc.edu/targetFinder/) using maximum stringency and NGG sites only (Table S6). *nAChRα3* Del sgRNA1 had one potential off-target and *nAChRα3* Del sgRNA2, five.

| sgRNA | sgRNA sequence | Off-target sequence | Line Checked? Dα3 1.7 | Dα3 5.3 | Dα3 11.4 | Dα3 15.1 |
|---|---|---|---|---|---|---|
| nAChRα3 Del sgRNA 1 | GTCGCCGCAA ACGCACACGA | GTtGgCcCAA ACGCAgACGA | Yes | No | No | No |
| nAChRα3 Del sgRNA 2 | GCCCATCGAT ATAAAGCTAT | attttTCtAT ATAAAGCTAT | Yes | No | No | No |
| | | GgaaAataAT ATAAAGCTAT | Yes | Yes | Yes | Yes |
| | | tatgtagtAT ATAAAGCTAT | Yes | No | No | No |
| | | GCCtAgatAT ATgAAGCTAT | Yes | No | No | No |
| | | taCCATaaAT ATAcAGCTAT | Yes | No | No | No |

**Table S6: List of the off-targets identified for the set of sgRNA used to generate deletions in *nAChRα3*.**

Off targets were tested through PCR and Sanger-sequenced for confirmation in an incomplete factorial design whereby all off-target regions were checked for strain 1.7, and off-target 3 was tested for all strains. The sequenced regions did not show the presence of a deletion using the primer set reported in Table S7.

| sgRNA | Off-Target sequence | Primer used | Primer sequence |
|---|---|---|---|
| nAChRα3 Del sgRNA 1 | GTtGgCcCAAACGCAgACGA | nAChRα3 Del OT1 Check F | GACACTGCCACCACCACAC |
| | | nAChRα3 Del OT1 Check R | TGGCGGGCAAAGTCATAACT |
| nAChRα3 Del sgRNA 2 | attttTCtATATAAAGCTAT | nAChRα3 Del OT2 Check F | CAGTTTCCTACACCTCGCGT |
| | | nAChRα3 Del OT2 Check R | GGTCGAGGTTTCCTTTTGAGC |
| | GgaaAataATATAAAGCTAT | nAChRα3 Del OT3 Check F | GCCAAACAATAACAGACGGCA |
| | | nAChRα3 Del OT3 Check R | TTTGCCGCTCAAAGTTTGGC |
| | tatgtagtATATAAAGCTAT | nAChRα3 Del OT4 Check F | ACTCCCTTTTGCCCCGTTAG |
| | | nAChRα3 Del OT4 Check R | TTCGCATGGCTCTGACCTTT |
| | GCCtAgatATATgAAGCTAT | nAChRα3 Del OT5 Check F | TGGCATCTAAACAAATCAGCCC |
| | | nAChRα3 Del OT5 Check R | CGCTACACCCACACACTAGT |
| | taCCATaaATATAcAGCTAT | nAChRα3 Del OT6 Check F | TGAGGGCAAAGGACTGTTACT |
| | | nAChRα3 Del OT6 Check R | GGGTGCCAAAAGAAGGGGAA |

**Table S7: List of primers used for off-target checking**

*4.2 Selection of candidate genes for functional validation*

A more comprehensive candidate gene list was selected based on the excess of significantly associated polymorphism within 2.5kbp of a gene coding sequence in at least two GWAS or present in 300bp segments with significant copy-number association, including 745 genes in total (Supplementary Data). 232 of these are expressed in adults, the stage tested in this study, and 205 are expressed in the adult head. Altough there was no functional annotation for 226 genes which limits the significance of the findings, 110 could be mapped to KEGG or Reactome pathways resulting in 59 connected genes showing enrichments for axon guidance and ensheathment and drug metabolism (Fig. S4, Monte Carlo simulation, $P<0.005$). Mutants with null alleles for *Ank* showed increased susceptibility while deletion for *UGT86Dj-h* and *Pde9* showed opposite temperature-dependent susceptibility (Fig S5).

A set of genes testing the polygenic basis of resistance was selected from the pathway and functional interaction network (Fig. S4) to cover a diverse subnetworks enriched for particular ontologies. *Ankyrin* was chosen to represent axon guidance and signal transduction based on a previous association study for SpinosAD resistance (Byrne, personal communication); *UDP-Glucose Transferase 86D-i* and *UDP-Glucose Transferase 86D-j* were chosen to represent drug metabolism based on a previous association with nicotine resistance; Phosphodiesterase 9 was chosen to represent purine metabolism based on a previous association with Chlorantraniliprole (Green et al., *in prep*).

Further three non-candidate control genes S*ol narae*, *Olfactory receptor 65a* and *α-Esterase 7* were used as random negative control genes and did not show any significant difference between MiMIC mutants and wildtype.

**Figure S4: Network of gene interactions significantly enriched for candidate genes conferring imidacloprid resistance.**

*(Figure S4 on next page)*

*4.2 Knock-out mutations and crossing design*

The source of alleles and the crossing design for the tests are reported in Table S6. For *Prm*, a Mi{MIC} knock-out mutant line carrying a heterozygous non-sense insertion in the first intron of the gene was ordered (Table S6) and backcrossed to the original y[1]w[*] line used for the Mi{MIC} insertion provided by the Bellen laboratory( http://flypush.imgen.bcm.tmc.edu/pscreen/about.html) . These were used to obtain heterozygous Prm mutants (selection of wildtypes and not *sb)* comparable to y[1]w[*] for control. For *nAChRα3*, homozygous knock-out mutants were generated using a CRISPR-CAS9 system. Homozygous *nachrα3* mutants carrying a {pActin-CAS9} transgene were directly compared to y[1]w[*]; {pActin-CAS9}. For Ank, a MiMIC knock-out mutant line carrying a homozygous non-sense insertion in the first intron of the gene was ordered. For *UGT86Di* and *UGT86Dj* two deletion lines were reciprocally crossed to generate a homozygous deletion of *Ugt86Dc-d-i-j* (selected to be wildtype and not sb) and compared to the deletion lines backcrossed to w[1118] (selected to be wildtype and not sb). For *Pde9*, two deletion lines were reciprocally crossed to generate a homozygous deletion restricted to *Pde9* (selection of wildtypes and not bar-eyed) and compared to the deletion lines backcrossed to w[1118] (selection of wildtype and not bar-eyed).

| Gene Affected | Stock Center id of allele donor | Full genotype of allele donor | Genotype tested | Control |
|---|---|---|---|---|
| *Prm* | 56623 | y[1] w[*]; Mi{y[+mDint2]=MIC}Prm[MI11769]/TM3, Sb[1] Ser[1] | y[1] w[*]; Mi{y[+mDint2]=MIC}Prm[MI11769]/+ and y[1] w[*]; +/Mi{y[+mDint2]=MIC}Prm[MI11769] | y[1] w[*] |
| *nACHRα3* | NA | y[1] w[*]; {pActin-CAS9}; nACHRα3[KO-1.7] | y[1] P(act5c-cas9, w+) M(3xP3-RFP.attP)ZH-2A w[*]; nACHRα3[KO-1.7] | y[1] P(act5c-cas9, w+) M(3xP3-RFP.attP)ZH-2A w[*] |
| *nACHRα3* | NA | y[1] w[*]; {pActin-CAS9}; nACHRα3[KO-5.3] | y[1] P(act5c-cas9, w+) M(3xP3-RFP.attP)ZH-2A w[*]; nACHRα3[KO-5.3] | y[1] P(act5c-cas9, w+) M(3xP3-RFP.attP)ZH-2A w[*] |
| *nACHRα3* | NA | y[1] w[*]; {pActin-CAS9}; nACHRα3[KO-11.4] | y[1] P(act5c-cas9, w+) M(3xP3-RFP.attP)ZH-2A w[*]; nACHRα3[KO-11.4] | y[1] P(act5c-cas9, w+) M(3xP3-RFP.attP)ZH-2A w[*] |
| *nACHRα3* | NA | y[1] w[*]; {pActin-CAS9}; nACHRα3[KO-15.1] | y[1] P(act5c-cas9, w+) M(3xP3-RFP.attP)ZH-2A w[*]; nACHRα3[KO-15.1] | y[1] P(act5c-cas9, w+) M(3xP3-RFP.attP)ZH-2A w[*] |
| *UGT86Di-j* | 9083 | w[1118]; Df(3R)ED5506, P{w[+mW.Scer\FRT.hs3]=3'.RS5+3.3'}ED5506/TM6C, cu[1] Sb[1] | w[1118]; Df(3R)ED5506, P{w[+mW.Scer\FRT.hs3]=3'.RS5+3.3'}ED5506/Df(3R)Exel7306 and w[1118]; Df(3R)Exel7306/Df(3R)ED5506, P{w[+mW.Scer\FRT.hs3]=3'.RS5+3.3'}ED5506 | w[1118]; Df(3R)ED5506, P{w[+mW.Scer\FRT.hs3]=3'.RS5+3.3'}ED5506/+ and w[1118]; Df(3R)Exel7306/+ |
| *UGT86Di-j* | 7957 | w[1118]; Df(3R)Exel7306/TM6B, Tb[1] | | |
| *Pde9* | 29028 | Df(1)BSC851, w[1118]/FM7h/Dp(2;Y)G, P{w[+mC]=hs-hid}Y | Df(1)BSC851/Df(1)BSC545, w[1118] and Df(1)BSC545/Df(1)BSC851, w[1118] | Df(1)BSC851/+, w[1118] and Df(1)BSC545/+, w[1118] |
| *Pde9* | 25073 | Df(1)BSC545, w[1118]/FM7h/Dp(2;Y)G, P{w[+mC]=hs-hid}Y | | |
| *Ank* | 60831 | y[1]; Mi{y[+mDint2]=MIC}Ank[MI10114] | y[1]; Mi{y[+mDint2]=MIC}Ank[MI10114] | y[1] w[*] |

**Table S7:** Mutant allele origin and crossing design used for the functional tests

*4.3 Analysis*

**Figure S5: Mean longevity +/- standard error of flies exposed to imidacloprid relative to control** (y-axis) **for wild type and mutants in candidate genes.** The top-right box in each panel is a scatter-boxplot recapitulating the difference in genetic effect between mutant and wildtype for each gene and temperature comparison in the same unit of relative longevity with respect to control; the thick shows the mean effect, the box shows the interquartile distribution and the whiskers, the 5%- and 95%-tile distribution. Test for *Ank* are shown in panel A and B, *UGT86Di-j* in panel C and D and *Pde9* on panel E and F. Tests performed at 20°C are reported on panels A, C and E; tests performed at 30°C are reported on panels B, D and F**.** **: P-value<0.01 (t-test).

*(Figure S5 on next page)*
